# An atomistic model of myosin interacting heads motif dynamics and their modification by 2’-deoxy-ADP

**DOI:** 10.1101/2024.06.06.597809

**Authors:** Matthew Carter Childers, Michael A. Geeves, Michael Regnier

**Affiliations:** Department of Bioengineering, School of Medicine, University of Washington, Seattle, WA, USA; Department of Biosciences, University of Kent, Canterbury, Kent CT2 7NJ, UK

**Keywords:** Myosin motor, interacting heads motif (IHM), super-relaxed state (SRX), molecular dynamics, muscle regulation, muscle structure and function

## Abstract

The contraction of striated muscle is driven by cycling myosin motor proteins embedded within the thick filaments of sarcomeres. In addition to cross-bridge cycling with actin, these myosin proteins can enter an inactive, sequestered state in which the globular S1 heads rest along the thick filament surface and are unable to perform motor activities. Structurally, this state is called the interacting heads motif (IHM) and is a critical conformational state of myosin that regulates muscle contractility and energy expenditure. Structural perturbation of the sequestered state via missense mutations can pathologically disrupt the mechanical performance of muscle tissue. Thus, the IHM state has become a target for therapeutic intervention. An ATP analogue called 2’-deoxy-ATP (dATP) is a potent myosin activator which destabilizes the IHM. Here we use molecular dynamics simulations to study the molecular mechanisms by which dATP modifies the structure and dynamics of myosin in a sequestered state. Simulations with IHM containing ADP.Pi in both nucleotide binding pockets revealed residual dynamics in an otherwise ‘inactive’ and ‘sequestered’ state of a motor protein. Replacement of ADP.Pi by dADP.Pi triggered a series of structural changes that modify the protein-protein interface that stabilizes the sequestered state, and changes to this interface were accompanied by allosteric changes in remote regions of the protein complex. A comparative analysis of these dynamics predicted new structural sites that may affect IHM stability.

## Introduction

Within striated muscle sarcomeres, the myosin motors of the thick filament bind to the actin strands of thin filaments and give rise to contraction (Figure 1A). (1, 2) In thick filaments, myosin motors form heterohexamers comprised of two myosin heavy chains (MHC), two essential light chains (ELC), and two regulatory light chains (RLC). (Figure 1B) Myosin heavy chains can be divided into the motor domain (Figure 1C) and a tail that adopts α-helical structure. The motor domain performs ATP hydrolysis, interacts with actin filaments, and triggers conformational changes associated with the powerstroke. The tail forms a coiled-coil structure and tails of multiple myosins intertwine and assemble with other proteins into thick filaments. The interacting heads motif (IHM) myosin state forms as the actin binding region of the ‘blocked head’ (BH) motor domain (Figure 1B, orange) interacts with the ‘free head’ (FH) via the ‘mesa’ (3). (Figure 1B, purple) This interaction is made possible through specific bending of a hinges in the myosin ‘neck’. (4, 5)

**Figure 1.**
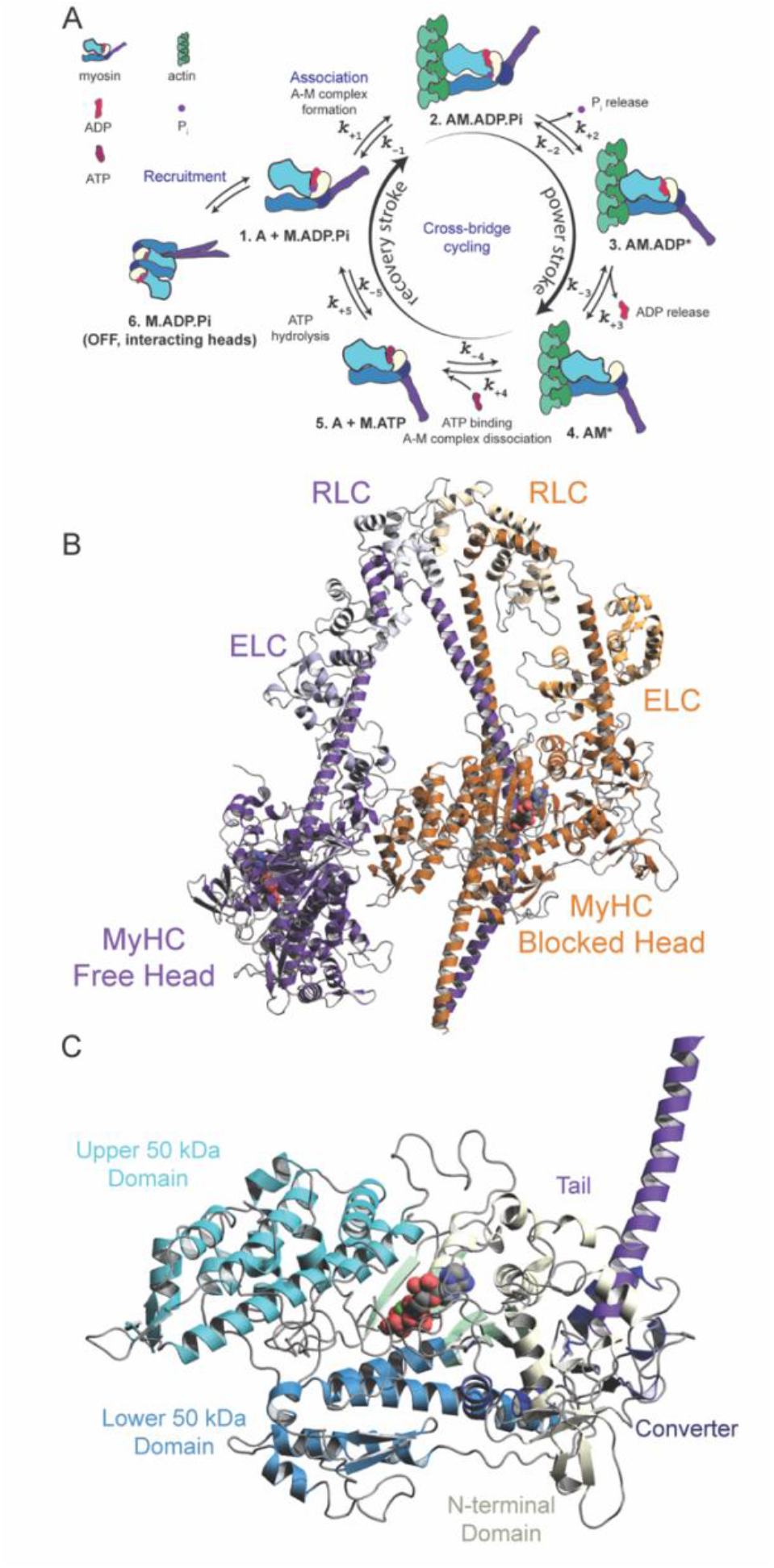
Schematic of the actin-myosin chemomechanical cycle. (A) Within sarcomeres, actin and myosin generate force through concerted interactions and conformational changes. This process, known as the cross-bridge cycle (steps 1 – 5), is driven by ATP hydrolysis. In addition to active conformations, myosin can access an inactive conformation in which the two motor domain heads of a striated muscle myosin II complex interact with one another (step 6). (B) Here we simulated a myosin IHM construct that contains a blocked head (orange) and free head (purple) along with their associated regulatory and essential light chains. Ligands in the nucleotide binding pockets are shown as spheres. (C) The motor domain of myosin can be divided into four large domains: the upper and lower 50 kDa domains form the actin binding and free head binding sites. The N-terminal domain and converter domain form a hinge site around which the tail bends during the powerstroke.

Force generation in muscle fundamentally depends on concerted interactions and dynamic conformational changes within and between myosin motors and actin filaments, in a process driven by ATP hydrolysis. (1) In addition to active, cycling conformations associated with actin binding and force generation (Figure 1A) (2), myosins can access inactive, non-cycling states with low ATP turnover. The low ATP turnover state has been associated with the structurally-defined IHM, where the two S1 myosin heads in the M.ADP.Pi state and sequestered into the folded back state. (6, 7) (Figure 1A) A folded back structure for the two head of myosin, that may be similar to the IHM state, is thought to be responsible for the slow rate of ATP turnover in stopped flow assays (8) and reduced ATP utilization (9) within the biochemically-defined super-relaxed (SRX) state of muscle tissue (10). For these reasons, the IHM has received attention for its modulation of muscle function (11–14), roles in pathological states (5, 15–17), and as a therapeutic target (8, 18–20).

The IHM is appreciated as a thick filament-based means of modifying myosin activity, myosin recruitment for sarcomere contractility, and muscle energy consumption. Muscle contractility is limited by the number of myosin heads available to interact with actin and consequently the force that can be generated by a thick filament. A rich set of structural and biochemical factors can modify myosin activity by affecting the stability of the IHM, including RLC phosphorylation, (21) myosin binding protein C phosphorylation, (22, 23) ionic strength, (19) and temperature (19, 24). Missense mutations in myosin can pathologically disrupt the natural equilibrium of free and sequestered myosins and lead to myopathies (3, 5, 15, 16). Small molecules have the potential to similarly modulate IHM stability (8, 19). In previous work, our group has studied the effects of the naturally occurring small molecule, 2’-deoxyATP (dATP), on the structure, dynamics, and function of myosin. We have reported that this ATP analogue not only affects myosin activity, (25–30) but also eliminates the slow ATP turnover rate as measured with stopped flow kinetics, (8) repositions myosin heads farther away from the thick backbone and closer to thin filaments, as observed with small angle X-ray diffraction of demembranated and intact muscle, and increases weak binding of myosin to thin filaments in resting muscle, as measured with sinusoidal stiffness analysis. (27, 30) Combined, these observations indicate that dATP destabilizes the IHM state.

The specific protein conformations and structural dynamics regulate IHM stability remain unclear. Increases in protein structural resolution using cryoEM have paved the way for studies of the IHM state. (6, 31–36) Grinzato *et al*. recently produced a high-resolution (3.6 Å) structure of IHM state human β-cardiac myosin. (37) Dutta *et al*. recently obtained structures of mavacamten-stabilized human ventricular C-zone thick filaments that include IHM state β-cardiac myosin (6.4 Å resolution) from human cardiac ventricular tissue. (38) Contemporaneously, Tamborrini *et al*. obtained structures of mavacamten-stabilized mouse C-zone thick filaments (18 Å resolution) as well as thin filaments using mouse cardiac ventricular myofibrils (39). However, these studies have examined static IHM conformations in static environments and the thick filament structures were obtained in the presence of mavacamten, which stabilizes an otherwise dynamic and heterogeneous environment. Comparatively few studies have examined the structures and dynamics of myosin under conditions that destabilize the IHM. Here, we utilize microsecond scale molecular dynamics (MD) simulations to investigate the intrinsic dynamics of the IHM and to probe the molecular mechanisms by which dATP destabilizes the IHM. Our study is comprised of two simulated IHM myosin systems; each system contains two copies of human β-cardiac myosin, essential light chain, and regulatory light chain along with solvating water molecules that amount to ∼600k atoms, and simulations were performed using Anton 2 (40). In the reference system (myosin + ADP.Pi) ADP, inorganic phosphate (henceforth Pi), and Mg^2+^ are bound in both nucleotide binding pockets. In the test system (myosin + dADP.Pi), both ADP.Pi molecules were replaced by dADP.Pi. We find that the dADP nucleotides were more dynamic than their ADP counterparts and formed distinct interactions with myosin. Altered myosin:nucleotide contacts propagated throughout the complex and modulated of residue-residue interactions at the BH – FH interface and contemporaneously altered structures and dynamics of the tail. Our simulations contribute to the growing body of work highlighting the structural effects of dADP.Pi on myosin structure/function.

## Results

### Dynamics within the IHM state

To assess the effects of dATP, we evaluated large-scale conformational changes observed during MD. We calculated the C_α_ root-mean-square deviations (RMSD) of the simulations referenced against the equilibrated structures. 13 different RMSD calculations were performed, each associated with unique subsets of protein C_α_ atoms (Table 1). For 10 out of these 13 calculations, the myosin+dADP.Pi simulation experienced greater deviations from the reference structure than the myosin+ADP.Pi simulations (Table 1). The myosin+dADP.Pi simulation experienced greater structural deviations from the equilibrated conformation, but the extent of structural deviation varied among the different structural regions.

**Table 1.**
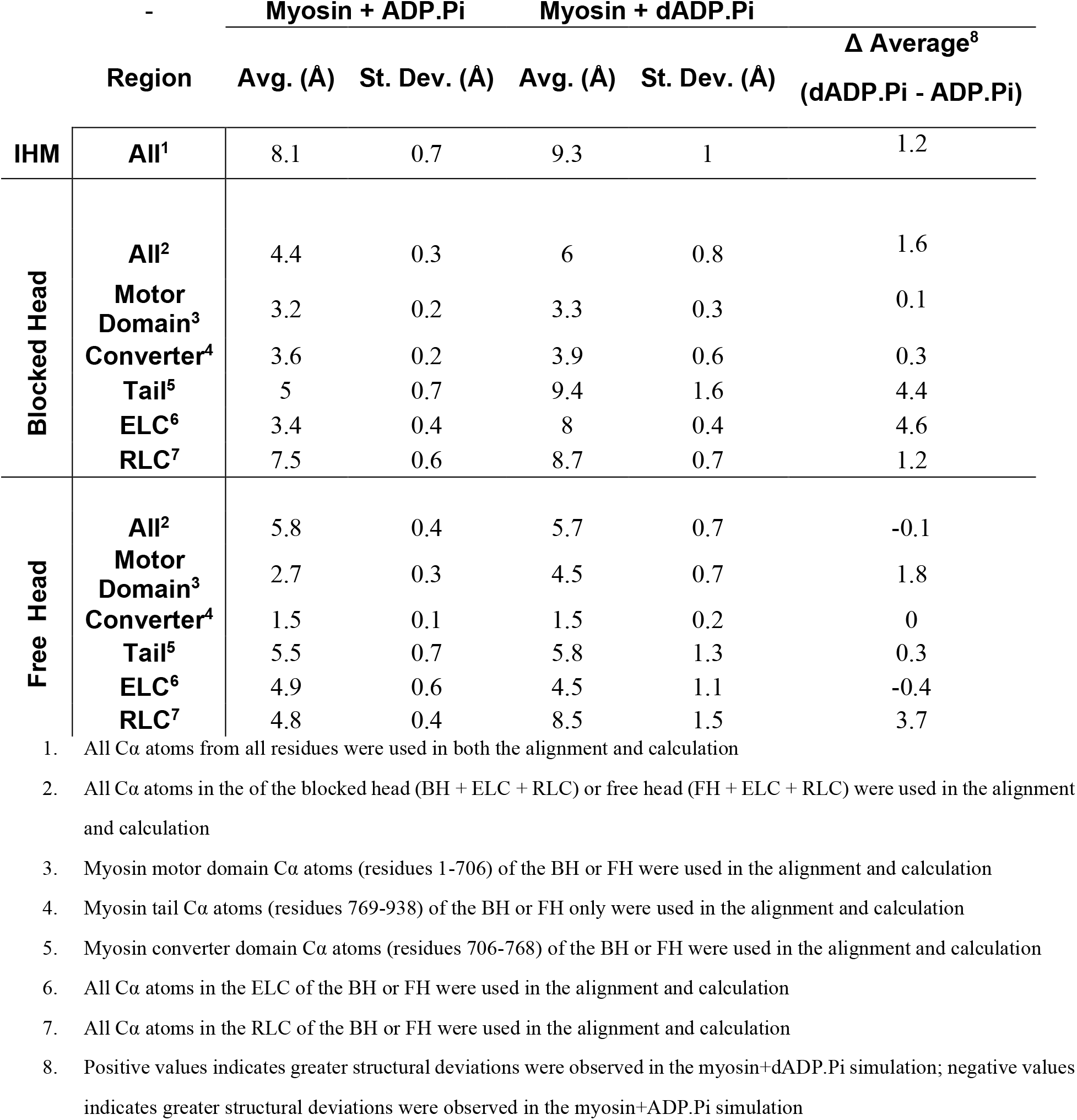
C_α_ RMSD of select myosin regions highlights that the dADP.Pi simulations experienced greater average structural deviations from the equilibrated starting conformation.

The simulated IHM systems contain many flexible components: we used conformational clustering to identify major structural changes related to IHM destabilization. To assess structural diversity within the simulations, we re-aligned the trajectories to a set of ‘minimally dynamic residues’ identified via the *mdlovofit* algorithm.(41–43) This set is comprised of 1,482 residues in the motor domains and motor domain-binding regions of the tail. After trajectory alignment, RMSD and RMSF values were calculated for all C_α_ atoms, but only the *mdlovofit* selected residues were used to perform the alignments (Figure 2A-C). As was observed in the first alignment, the myosin+dADP.Pi system was more dynamic and sampled conformations further away from the equilibrated reference structure than the myosin+ADP.Pi system (Figure 2A). Also, residues in the myosin+dADP.Pi simulation experienced greater atomic fluctuations relative to the myosin+ADP.Pi simulation (Figure 2B-C). In both simulations, residues in the FH had higher atomic fluctuations than in the BH, and residues in the light chain-binding portions of the tail, the ELC, and RLC had the greatest overall atomic fluctuations (Figure 2B-C). To visualize major structural changes in the IHM simulations and assess conformational heterogeneity within each simulation, we clustered the MD trajectories using the *NMRclust* algorithm (44) implemented in *UCSF Chimera* (45). The simulations were clustered on the C_α_ atoms of residues 1-892 of both the BH and FH and trajectories were subsampled every 12 ns. The pairwise C_α_ RMSDs between each cluster representative for the myosin+ADP.Pi and myosin+dADP.Pi simulations (Figure 2D) show that the myosin+dADP.Pi simulation had greater conformational heterogeneity within the myosin+dADP.Pi simulation. The top 5 most populous clusters for each simulation shows that structural changes in the light chain binding regions of the tail and the BH – FH interface contributed most to the structural differences between the simulations (Figure 2E). This coarse-grain analysis of the IHM dynamics demonstrates that in solution the IHM is a dynamic protein complex that samples multiple conformations and that increased structural changes due to the replacement of ADP.Pi by dADP.Pi are observable on microsecond timescales.

**Figure 2.**
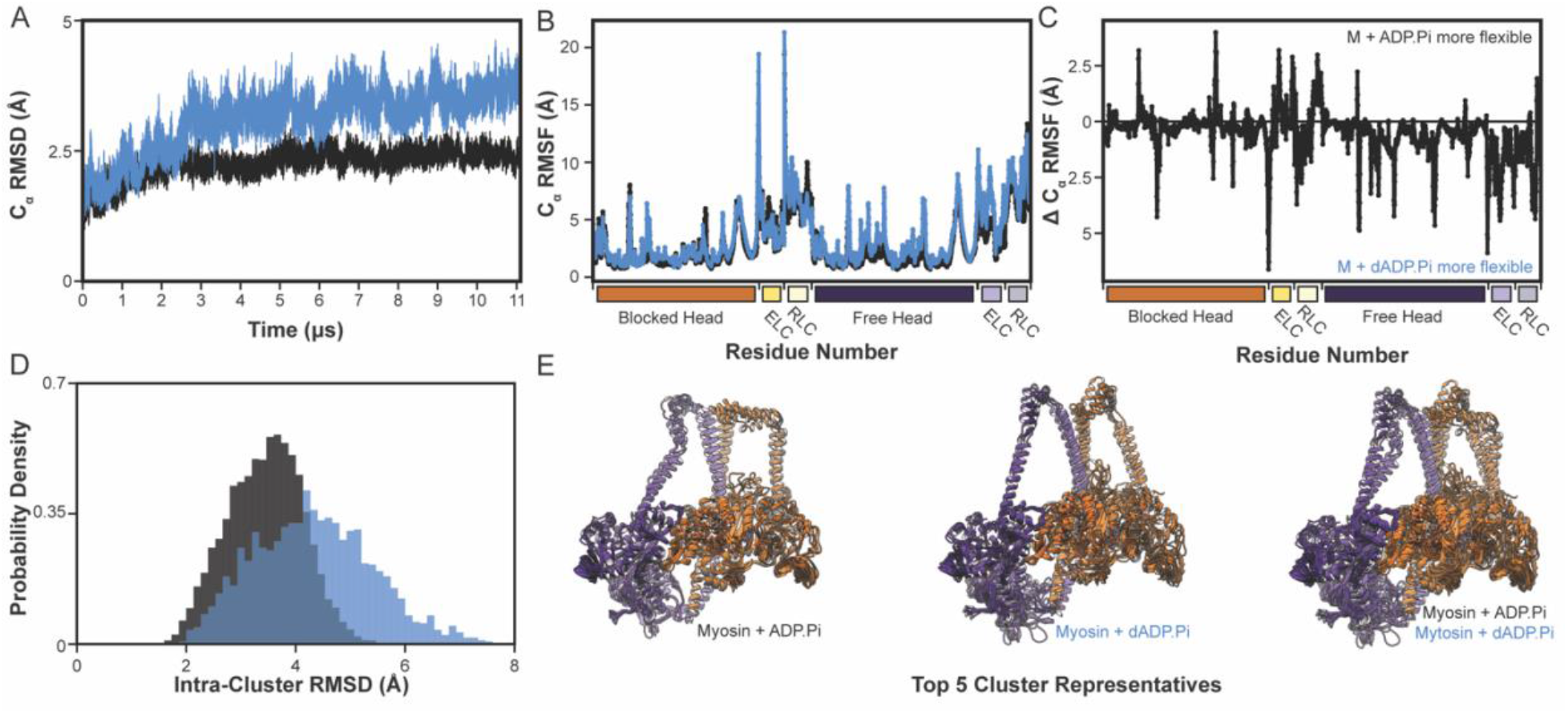
Overall dynamics of IHM myosin. (A) C_α_ RMSD of the IHM myosin + ADP.Pi (black) and IHM myosin + dADP.Pi (blue) simulations as a function of simulation time. RMSDs were calculated after alignment to stable residues identified by *mdlovofit*. (B) C_α_ RMSF of the IHM myosin + ADP.Pi (black) and dADP.Pi (blue) simulations as a function of residue number. (C) The difference in C_α_ RMSF between the ADP.Pi and dADP.Pi-bound myosin simulations. Positive values indicate that residues were more flexible in the ADP.Pi simulation. (D) The distributions of the intra-cluster C_α_ RMSD values among the cluster representatives for the ADP.Pi (black) and dADP.Pi (blue) simulations. (E) The cluster representatives for the top 5 most populous clusters in the ADP.Pi (left) and dADP.Pi (center) simulations are shown and aligned on the *mdlovofit* selected residues. The image on the right superimposes the cluster representatives from both simulations.

### dADP.Pi formed distinct interactions with myosin motor domains

To identify the molecular origins of the dynamic differences between myosin+ADP.Pi and myosin+dADP.Pi, we compared the BH nucleotide binding pockets when bound to the two different nucleotides. Nucleotide dynamics were evaluated by monitoring 6 dihedral angles that describe (d)ADP structure using internal coordinates, as previously described (28). The 6 calculated dihedral angles were defined as: θ_1_ (atoms: O1B-PB-O3A-PA), θ_2_ (PB-O3A-PA-O5′), θ_3_ (O3A-PA-O5′-C5′), θ_4_ (PA-O5′-C5′-C4′), θ_5_ (O5′-C5′-C4′-O4′), and θ_6_ (O4′-C1′-N9-C8) (Figure 3A). Ramachandran-style dihedral angle maps (θ_1_ vs θ_2_, θ_3_ vs θ_4_, and θ_5_ vs θ_6_) describe the relative orientation of the phosphate groups, the sugar ring, and the purine ring. Nucleotide conformation was strongly related to the interactions formed with residues in the nucleotide binding pocket of myosin (Figure 3B). The dihedral maps show that ADP.Pi and dADP.Pi were dynamically and structurally distinct and that the dADP nucleotides in both the blocked and FHs sampled a greater degree of conformational space and had greater conformational entropy (Figure 3C). These results mirror prior studies of dATP/dADP demonstrating that dADP (Figure 3E) is an intrinsically more dynamic nucleotide than ADP (Figure 3F) when bound to myosin (25, 28).

**Figure 3.**
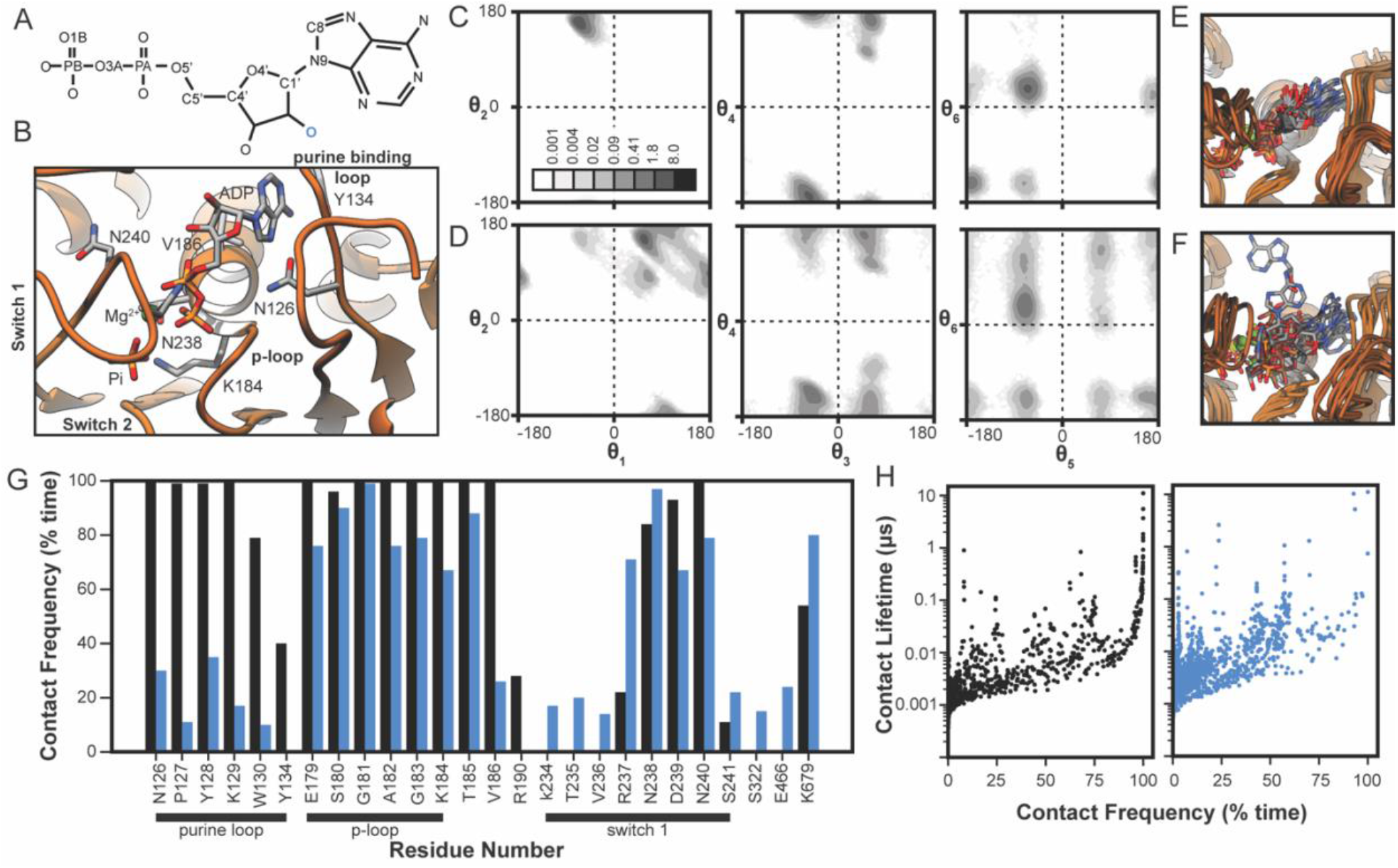
dADP was more dynamic than ADP. (A) The structure of an ADP molecule is shown. The names of atoms that define the dihedral angles are provided in the structure and the 2’ oxygen atom that is absent in dADP is colored blue. (B) Snapshot of the equilibrated conformation of the nucleotide binding pocket of the blocked head identifies functional residues in the purine binding loop, phosphate binding loop, and switch 1 that participate in interactions with ADP. Ramachandran-like dihedral angle maps of 6 angles that collectively describe the nucleotide conformations of ADP (C) and dADP (D) within the nucleotide pocket of the blocked head. (θ_1_, θ_2_) describes the relative orientation of the terminal phosphate groups (left); (θ_3_, θ_4_) describes the orientation of the sugar relative to the alpha phosphate (center); (θ_5_, θ_6_) describes the orientation of the purine ring relative to the sugar ring (right). Dihedral angle data were allocated to 5º x 5º bins and shaded according to the number of conformers allocated to each bin. Inset color scale indicates the percentage of the simulated ensemble allocated to each 5º x 5 º bin. (G) The percent of simulation time for which different myosin residues (x-axis) interacted with ADP (black) and dADP (blue) are shown. Calculations were performed at the residue level: a nucleotide-residue pair was considered ‘in contact’ if at least one pair of heavy atoms had a distance < 5 Å. Nucleotide-myosin interactions were nucleotide-dependent. (H) Atom-atom contact lifetimes as a function of contact frequency are shown for myosin:ADP (black, left) and myosin:dADP (blue, right) interactions within the blocked head. Calculations were performed at the atom level: each point corresponds to a contact between a unique pair of heavy atoms.

The distinct dynamics of ADP and dADP in the BH resulted in distinct interactions between the nucleotides and myosin residues within the nucleotide binding pocket. We measured the percent simulation time for which the nucleotides interacted with a myosin residue (Figure 3G). In the myosin+ADP.Pi simulation, the nucleotide retained its starting conformation that has been observed in structure of pre-powerstroke (1. M.ADP.Pi) state myosin. The purine ring of the nucleotide interacted with the purine binding loop (residues 126-134), the phosphate groups remained surrounded by the phosphate binding loop (residues 178-184), and the sugar ring formed transient interactions with residues in switch 1 (residues 232-244). In contrast, the more dynamic dADP formed fewer interactions with purine loop residues, fewer interactions with phosphate binding loop residues, and altered interactions with switch 1(Figure 3G). Finally, we examined the average duration of a nucleotide:myosin contacts at the atomic level (Figure 3H). We found that dADP participated in fewer long-lived (i.e. > 75% contact frequency) interactions and that the average atom-atom contact lifetime was decreased for myosin+dADP.Pi (Figure 3H). Similar behavior was observed for the nucleotides in the FH.

### Altered nucleotide-myosin interactions propagated throughout the IHM structure

To determine the impact of the more dynamic dADP.Pi on IHM dynamics, we next analyzed residue-residue contacts in the IHM. For each MD snapshot, two residues were considered in contact if at least one pair of heavy atoms between them were within 5Å of one another. We compared the residue-residue contacts formed in the simulations and only considered contacts which were present for at least 10% of time in either simulation. Contacts were considered as altered by dADP.Pi if there was at least a 30% difference in the percentage of simulation time that the contact was present in the two simulations. With this conservative definition, of the ∼29,000 observed residue interactions, ∼3,500 contact pairs differed by at least 30% between the myosin+ADP.Pi and myosin+dADP.Pi simulations. We mapped the altered interactions onto the equilibrated starting structure of the myosin+ADP.Pi simulation (Figure 4A). The altered contacts clustered in three regions of the IHM: the upper 50 kDa domains of both the BH and FH, the BH – FH interface, and the interfaces between the myosin heavy chains and light chains (Figure 4A). The pattern of contacts altered by dADP.Pi suggest that changes in the nucleotide binding pocket allosterically propagated through the upper 50 kDa domain of the BH and result in altered interactions made by nearby loops (residues 275-282, 311-324), the O-helix (residues 416-448), and ultimately the functional loops: loop 2 (residues 622-645), loop 4 (residues 361-377), and the cardiomyopathy loop (residues 402-416) (Figure 4A).

**Figure 4.**
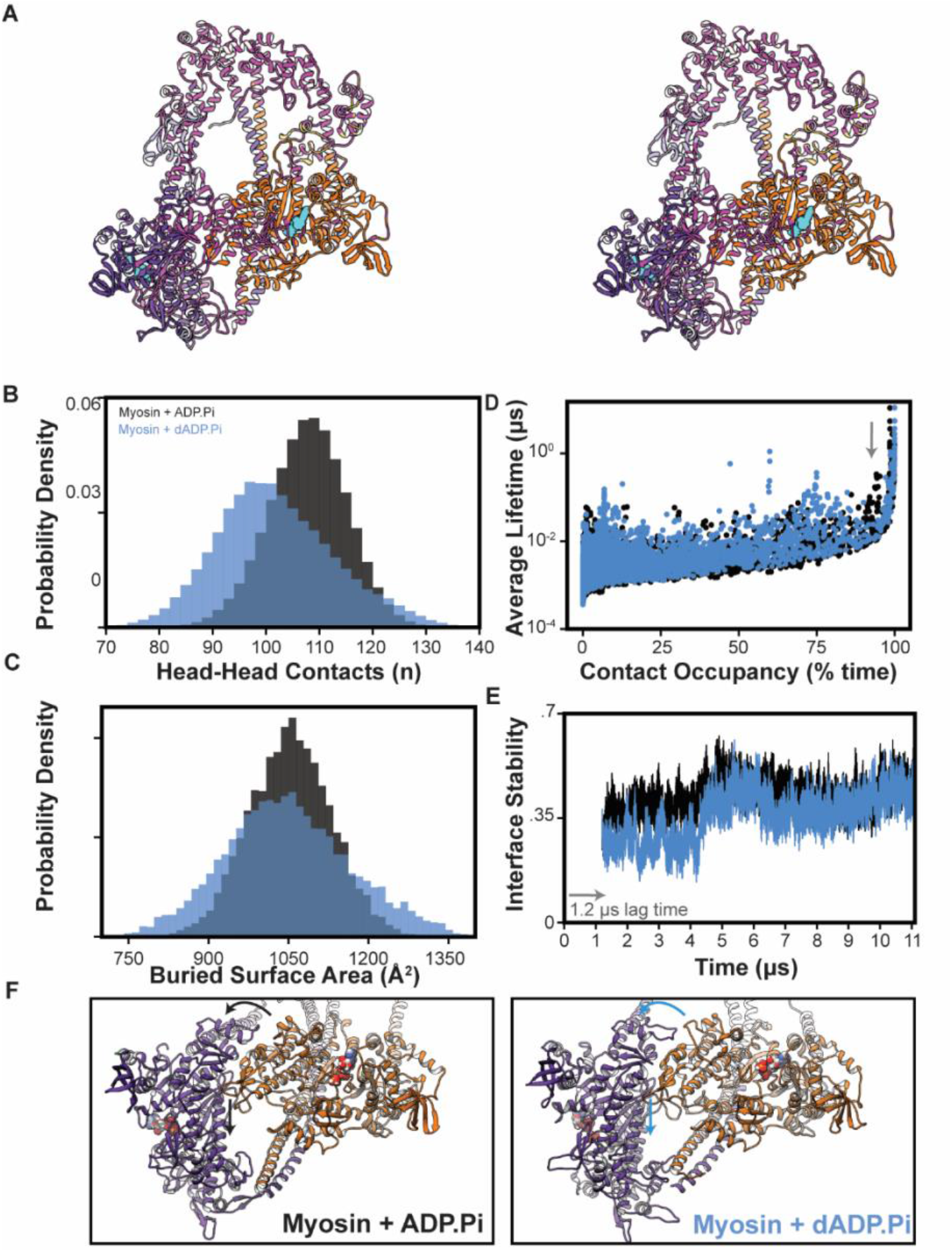
Altered nucleotide dynamics led to altered conformations of the motor domain. (A) Stereo image of the IHM depicting structural changes between the ADP.Pi and dADP.Pi simulations. Residues that had altered contacts are shown in magenta. (B) The total number of interacting residue pairs between the blocked head and free head were monitored over the course of the simulation. (C) The surface area of the interface between the blocked head and free head was variable during the simulations had was more variable in the myosin+dADP.Pi simulation. (D) Contact lifetimes versus contact occupancies for the atom-atom interactions between the blocked head and free head in the ADP.Pi (black) and dADP.Pi (blue) simulations. (E) Jaccard index calculated for the set of atoms participating in the blocked head – free head interface as a function of time with a 1.2 microsecond lag. (F) Changes to the blocked head – free head interface are highlighted on structural snapshots taken at the end of the simulations. Relative to the ADP.Pi simulation, the motor and ELC of the free head bent away from S2 and the blocked head.

### Conformational heterogeneity at the blocked head – free head interface

In the IHM state, the two motor domains form an evolutionarily conserved interface in which the mesa (3, 34) of the BH docks against the FH and the coiled-coil S2 tail. Changes in the internal structure of the upper 50 kDa domain (Figure 4A) of the BH led to modulation of the BH – FH interface. To assess the stability of the BH – FH interface, we tracked the interacting residue-residue pairs between the BH and FH over time. Independent of the bound nucleotide, we found that the interface was intrinsically dynamic. The number and composition of interacting residue pairs was variable in the myosin+ADP.Pi simulation. BH – FH interaction was not a fixed interface, but like other protein complexes the interface was fluid and dynamic. (46) In a time average, myosin+dADP.Pi had 7% fewer BH – FH interactions than myosin+ADP.Pi, though there were conformations observed with increased and decreased BH – FH interactions (Figure 4B). The surface area of the buried BH – FH interface (Figure 4C) was similar in both simulations, though there was more heterogeneity in myosin+dADP.Pi. For either case, the conformations explored by myosin+dADP.Pi were more heterogeneous. The average contact lifetimes for atom-atom pairs in the two simulations (Figure 4D) and observed that the myosin+dADP.Pi simulation had slightly fewer long-lived inter-head contacts than the myosin+ADP.Pi simulation (Figure 4D) To evaluate the rate of change of the BH – FH interface, we calculated the Jaccard Index of residue pairs in the BH – FH interface for every frame *i* versus frame *i-* 10,000 (a 1.2 microsecond lag, Figure 4E). There were low values of the Jaccard index for both simulations during the first ∼4 microseconds of the trajectories, though the value for myosin+dADP.Pi was lower, indicating a greater rate of change. By the end of the simulations, the interfaces for both models stabilized to a similar degree, but still retained some dynamic fluidity, as has been reported in studies of other protein interfaces (46). Changes in the structure of the upper 50 kDa domain associated with the replacement of ADP.Pi by dADP.Pi ultimately led to a distinct interface formed between the BH and FH (Figure 4F).

### Conformational heterogeneity within the light chain binding domain

The second site of major conformational change identified by our contact analysis were the regions of myosin that interacted with the light chains. In addition, there are several residues in the pliant region of the tail that form hinge sites that enable the heads to adopt various orientations and likely contribute to the formation of the IHM. To analyze the tail conformations in our simulations, we measured four interhelical angles formed between the flexible joints of the motor domain, lever arm, and S2 tail (Figure 5A). Angle 1 was defined by residues 699-706 and 770-783; angle 2 by 786-800 and 812-822, angle 3 by 812-822 and 831-838, and angle 4 by 831-383 and 845-872. After the equilibration phase that occurred at the beginning of the simulations, the hinge sites sampled conformations around an average value. For most angles the average values were different in the myosin+ADP.Pi and myosin+dADP.Pi simulations (Figure 5B). There were also differences in the average angles sampled by the BH and FH heavy chains (Figure 5B). Certain pairs of interhelical angles had correlated motions. For example, there was a marked positive correlation between angle 2 of the FH and angle 3 of the BH in both simulations (Pearson’s R = 0.46 and 0.39, respectively) (Figure 5C-E). The distinct bending of the tails in the simulations was associated with a change in the orientation of the FH relative to the S2 tail and the BH (Figure 5F). In these simulations, the conformation and dynamics of the pliant region of the tail were nucleotide-sensitive and related to changes in the BH – FH interface. Thus, our model predicts the tail conformations adjust to accommodate alternate BH – FH orientations (or vice-versa).

**Figure 5.**
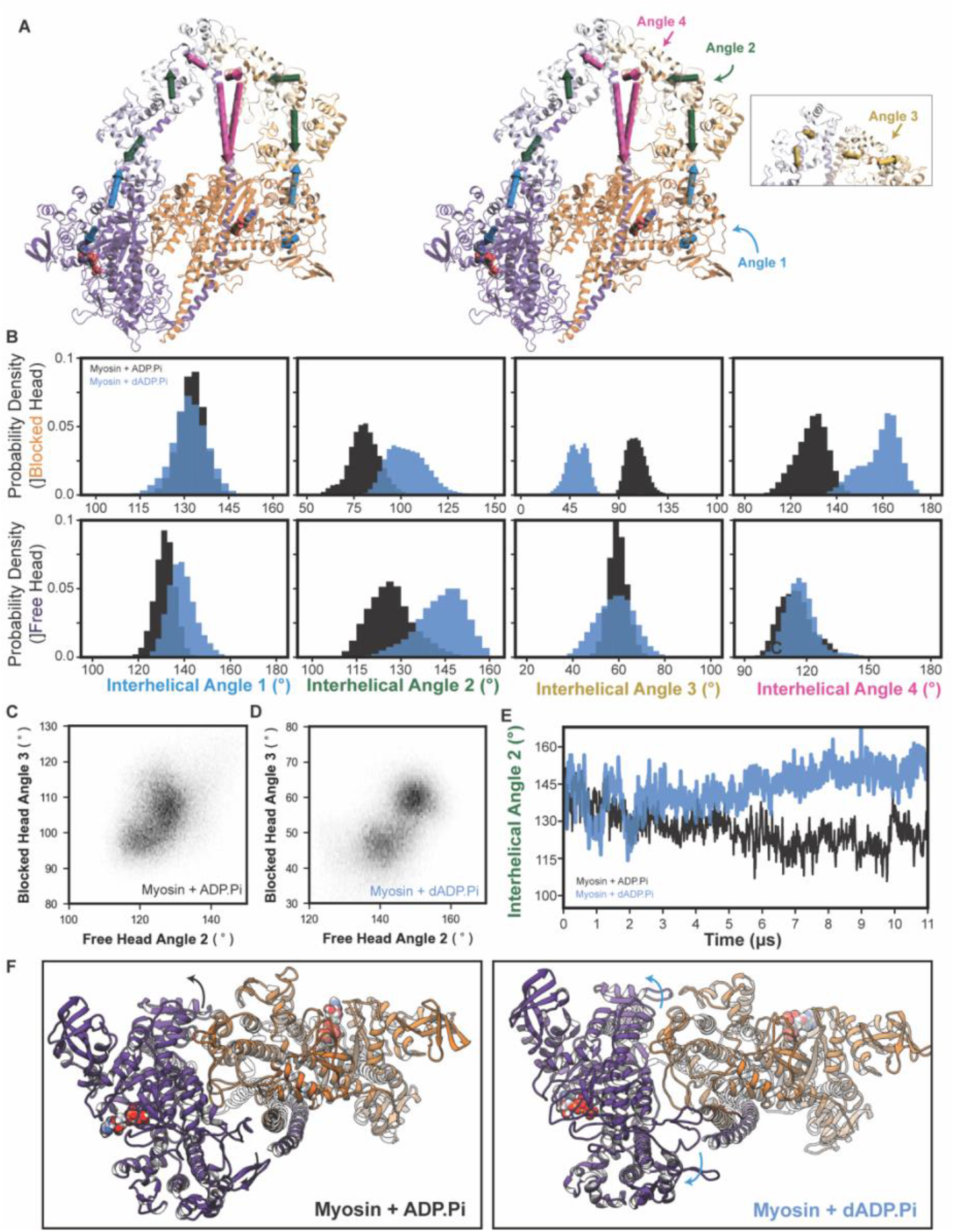
The tail of the free head adopted a distinct, ‘crimped’ conformation in the myosin+dADP.Pi simulations. Four interhelical angles (angles 1-4, blue, green, yellow, and magenta respectively) were analyzed over the course of the simulations. Panel (A) depicts a stereo image of the initial IHM structure with the four interhelical angles superimposed on the structure. (B) Distributions of the four interhelical angles calculated over the simulations are shown for the angles within the blocked head (top row) and free head (bottom row) for both the myosin+ADP.Pi (black) and myosin+dADP.Pi (blue) simulations. 2D histograms of the joint distribution of free head angle 2 and blocked head angle 3 for the ADP.Pi simulation (C) and dADP.Pi simulation (D) show a strong correlation between these angles. (E) A plot of the blocked head interhelical angle 2 as a function of time demonstrates a phase of rapid change at the beginning of the simulations followed by a period of angle stabilization. (F) As with changes to the blocked head – free head interface, changes in the tail angles were associated with altered orientation of the free head relative to S2 and the blocked head.

## Discussion

### The conformation and dynamics of the pliant region are correlated with the BH – FH interface

It has been previously shown that phosphorylation of the regulatory light chain (21) modifies the stability of the IHM and, further, that mutations in the pliant region of the tail of myosin can (de)stabilize the IHM (5). Our simulations provide insights into the structural mechanisms by which the conformation and dynamics of the tail/light chain modify IHM stability. First, our myosin+dADP.Pi simulation predicts that structural changes at the BH – FH interface are connected to corresponding changes in the conformations of S2 and the pliant region of the tail. We propose that flexibility in the tail of the BH is necessary for the heads to form distinct relative orientations and to ultimately detach from one another. The hinge sites C-terminal to the RLC binding site (angle 2 and 3, Figure 5) experienced the greatest changes in the two simulations and had high correlated motions, suggesting flexibility in this region is critical for native-like IHM stability. There are multiple HCM/DCM mutations in this region, including R783H, R787C, A797T, L811P, F834L, and N847 that may lead to myopathy by modifying IHM stability. The large-scale structural changes corresponding to reorganization of the head interface and being of the tails occurred over a 2-4 microsecond time period. This implies that the equilibration period for these IHM constructs is on the ∼2 microsecond time scale, highlighting the importance of access to long timescale simulations for investigating the structural and dynamic behavior of the IHM.

### The importance of the pliant region and dependence of IHM conformation and dynamics on environment

The simulations described here represent the longest simulations of human β-myosin in the IHM state ever performed at atomic resolution and in explicit solvent. Analysis of the myosin+ADP.Pi simulation alone yields powerful insights into the intrinsic dynamics of myosin in the IHM state. As with any molecular dynamics study, the simulations presented here suffer from the *accuracy* and *sampling problems* (48), and the results must be interpreted through these limitations. First, these simulations were initiated from homology models constructed using low-resolution (> 10 Å) cryoEM maps and did not begin from empirically observed structures (the *accuracy* problem). Grinzato *et al*. showed that initial homology models differed from empirical models with respect to the angles adopted by the tail and the residues participating in the BH – FH interface (37). A notable difference is in the conformation of the BH pliant region. In the pre-production portion of our study, we observed that the hinges in this region of the homology model transitioned to conformations similar to those in the empirical cryoEM model, though an exact correspondence was not observed. (Supplemental Movie 1) Second, these simulations have each probed 11 microseconds of IHM dynamics, which is extensive given contemporary MD capabilities but limited in biochemical contexts (the *sampling* problem). Given current structural data and simulations, it is not possible to assess the extent of sampling obtained in these simulations nor is it possible to assess whether the simulations have adopted a local conformational minimum or are *en route* to it (49).

### dADP.Pi modified the IHM conformation by modifying upper 50 kDa domain structure and the BH – FH interface

Despite dATP/dADP differing from ATP/ADP structurally by only a single atom, the two nucleotides have distinct effects on the stability of the inactive conformation of myosin, as well as on the kinetics of actively cycling myosin heads (8, 27, 30). Previously, we have observed nucleotide-dependent dynamics for multiple myosin sequences and in multiple chemomechanical states using diverse MD force fields and simulation engines. (25, 28–30) We have consistently observed that the two nucleotides sample distinct conformational ensembles, form distinct sets of interactions with the nucleotide binding pocket (particularly switch 1 and the purine binding loop), and each impact the organization of residues in the upper 50 kDa domain of myosin. Here, we used dADP as a tool to understand the factors that destabilize the IHM. We observed that replacement of ADP.Pi by dADP.Pi modified myosin structure via structural changes that propagated along an allosteric pathway passing through switch 1 and the upper 50 kDa domain. These results suggest that amino acid mutations within the upper 50 kDa domain have the potential to modify the upper 50 kDa domain structure, the presentation of charges and patterns of charged residues on the motor domain surface, and ultimately the stability of the IHM even if they do not directly participate in BH – FH interactions. Using their homology model, Alamo *et al*. identified mutations on the myosin mesa that could (de)stabilize the BH – FH interface.(15) Our simulations extend this list of pathologic mutation sites (47) to include residues that lie along the computationally identified allosteric pathway: N232H, D239N, R243H, Y283C, S291F, D309N, A381D, L427M, and V606M.

We speculate that the tail conformation observed in recent cryoEM conformations is a rigid conformer that may be stabilized by mavacamten, which is reported to bind to the same binding pocket as omecamtiv mecarbil and affects the lever arm position (50). Modeling the pliant region and particularly the interface between the RLCs is challenging and is a point of structural heterogeneity among cryoEM models of the IHM. The higher resolution of residues within the BH – FH interface coupled with the biomedical impact of disease associated mutations in the mesa have focused modeling efforts on the BH – FH interface (51–53). Many models of the pliant region and RLC have used molluscan RLC conformers as templates, and the correspondence of molluscan and mammalian light chain structures in unclear as molluscan RLCs can bind Ca^2+^ to regulate thick filament activity (54). Pliant region conformers have also been modeled using thick filament maps in which IHM motor domains form intermolecular interactions with neighboring IHMs, which may be heterogeneous (38, 39). Uncertainty around the pliant region and RLC conformation continues to impact structural and dynamical studies of the IHM. The Grinzato *et al*. model also indicated higher flexibility (lower resolution) within S2, the RLCs, and the light chain binding region of the tail. Flexibility within the pliant region may allow myosins to form the IHM in different thick filament geometries or to adopt distinct IHM conformers in different regions of the thick filament (55, 56). Flexibility among the hinge regions and heterogeneous interacts with other thick filament proteins may ultimately affect the rates of entry into and exit from the IHM. The simulations reported here indicate correlated motions among interhelical angles and a relationship between the conformations of the pliant regions and the BH – FH interface; however, the extent to which these simulations are ‘correct’ is unclear as the challenges associated with model building necessarily bias MD calculations. These findings should motivate increased attention to both the structure and dynamics of the pliant region and the RLC interface: higher resolution structural insights into this region are needed as the structural factors impacting IHM stability extend beyond the myosin mesa. In future studies, MD simulations may explore accessible conformations of the pliant tail in different structural, physical, and chemical environments.

## Conclusions

We propose that our simulations have captured structural changes that precede BH – FH dissociation and departure from the IHM conformation. Changes to IHM involved a three-part mechanism. First, replacement of ADP by dADP led to a series of structural changes that propagated through the upper 50 kDa domain. Next, changes in the upper 50 kDa domain altered the BH – FH interface. Finally, changes in the BH – FH interface were met with structural changes in the pliant region. This indicates that structural perturbations distant from the mesa (i.e. within the motor domain and the pliant region) may pathologically or therapeutically modify IHM stability. Despite several limitations, our study identifies new avenues for research into the structural and functional role of the IHM. In the context of the highly structured and dynamically-driven sarcomere, understanding protein structural ensembles is essential to model and engineer muscle behavior. The dynamic computational models utilized here indicate that the IHM is not a fixed, rigid state. Instead, the BH – FH interface and bent tails harbor intrinsic flexibility. These observations should inform ongoing and future experimental studies of HCM/DCM mutations and therapeutics that target the IHM.

## Methods

Two 11 microsecond-long molecular dynamics (MD) simulations of IHM state myosin (M.ADP.Pi) were performed with either ADP.Pi or dADP.Pi in the nucleotide binding pocket using *Anton 2* (40). In the ‘reference’ system (myosin + ADP.Pi), ADP, inorganic phosphate (P_i_), and Mg^2+^ were bound in nucleotide binding pockets of both the blocked and free head. In the ‘destabilizing conditions’ system (myosin + dADP.Pi), both ADP molecules were replaced by dADP. These systems were prepared for production runs using the *AMBER20* MD package. One simulation of each system was performed for ∼11 microseconds (μs) using the *Desmond MD* package. Detailed methods for system development, parameterization, simulation, and analysis may be found in the Supplemental Material.

## Acknowledgements

Molecular simulations were performed using Anton 2 computer time (allocation MCB210020P to MR), which was provided by the Pittsburg Supercomputing Center (PSC) through Grant R01GM116961 from the NIH. The Anton 2 machine at PSC was generously made available by D.E. Shaw Research. This work used the Extreme Science and Engineering Discovery Environment (XSEDE) resource COMET through allocation TG-MCB200100 to MR. XSEDE was supported by the National Science Foundation grant number ACI-1548562. Partial funding for MCC was provided by Award Number T32HL007828 from the National Heart, Lung, and Blood Institute. The content is solely the responsibility of the authors and does not necessarily represent the official view of the NHLBI or the NIH. This research was supported by the University of Washington Center for Translational Muscle Research (CTMR) via the National Institute of Arthritis and Musculoskeletal and Skin Diseases of the National Institutes of Health award number P30AR074990 and R01HL128368 to MR. We are grateful to Drs. William Lehman and Anne Houdusse for productive discussions.

