## Supporting Information for "An atomistic model of myosin interacting heads motif dynamics and their modification by 2’-deoxy-ADP"

### Supplemental Movie 1

The movie depicts an interpolated morph (i.e. non-physical atomic trajectory) between two conformations of the IHM used in this study: the initial and pre-production conformer conformers (chains colored as in Figure 1). The morph is superimposed on the static image of the Grinzato *et al.* cryoEM conformer (transparent, blue ribbons). This trajectory should not be interpreted as a realistic transition between two microstates; its intent is instead to highlight regions of structural change.

### Methods

Myosin+(d)ADP.Pi Model Building: Myosin heavy chain tails were truncated at residue 938. Then flexible loops were re-modeled over 3 successive rounds of loop modeling by *Modeller* (6, 7) followed by minimization in AMBER. The long ~40 residue N-terminal extension of the ELC was not modeled. The final systems includes all myosin heavy chain residues (1-938) with ADP, Pi, and $Mg^{2+}$ in the catalytic pockets of both motor domains, ELC residues (50-191), and RLC residues (2-159). A duplicate model (myosin+dADP.Pi) was generated in which the ADP nucleotides of both the blocked head and free head were replaced by dADP via removal of the 2' oxygen. His protonation states at pH 7.0 were predicted using *H++* using default parameters.(8)

Myosin+(d)ADP.Pi Simulation Preparation (myosin+ADP.Pi and myosin+dADP.Pi): The resulting systems were prepared for molecular dynamics simulation using the Amber 20 (9) simulation package and the ff14SB force field (10). Water molecules were treated with the TIP3P force field (11). Metal ions were modeled using the Li and Merz parameter set (12–14). ADP, dADP, and Pi (modeled as $H_2PO_4$ (15)) molecules were treated with the GAFF2 force field (16). Partial charges were derived from a restrained electrostatic potential (*resp*) fit to quantum mechanics calculations performed with ORCA (17). The SHAKE algorithm was used to constrain the motion of hydrogen-containing bonds (18). Long-range electrostatic interactions were calculated using the particle mesh Ewald (PME) method. Hydrogen atoms were modeled onto the initial structure using the *leap* module of *AMBER*, and each protein was solvated with explicit water molecules in a periodic, cubic box that extended at least 10 Å beyond any protein atom. $Na^+$ and $Cl^-$ counterions were added to neutralize the systems. Each system contained ~600,000 atoms. Each system was minimized in three stages. First, hydrogen atoms were minimized for 1000 steps in the presence of 100 kcal/mol restraints on all heavy atoms. Second, all solvent atoms were minimized for 1000 steps in the presence of 25 kcal/mol restraints on all protein atoms. Third, all atoms were minimized for 18000 steps in the presence of 25 kcal/mol restraints on all backbone heavy atoms (N, O, C, and $C_\alpha$ atoms). After minimization, systems were heated to 310 K using the NVT (constant number of particles, volume, and temperature) ensemble and in the presence of 25 kcal/mol restraints on backbone heavy atoms. Next, the systems were equilibrated over five successive stages using the NPT (constant number of particles, pressure, and temperature) ensemble. During the first four stages, the systems were equilibrated for 5 ns in the presence of 25 to 1 kcal/mol restraints on backbone heavy atoms. During the final equilibration stage, the systems were equilibrated for 50 ns in the absence of restraints. A 10 Å nonbonded cutoff was used for all preparation and production simulations.

Myosin+(d)ADP.Pi Molecular Dynamics Simulation: The equilibrated systems were then prepared for simulation on Anton 2 using the Desmond simulation package. Briefly, the equilibrated starting coordinates and parameter/topology files using *viparr-convert-prmtop*. Finally, ~11 μs long trajectories for each of the myosin+ADP.Pi and myosin+dADP.Pi systems were performed using the standard Anton 2 simulation protocols in the isobaric-isothermal ensemble and coordinates were saved every 120 picoseconds for analysis.

Structural and Dynamic Analyses: MD trajectories from Anton2 (myosin+(d)ADP.Pi) were analyzed at 120 ps granularity (~90000 snapshots) unless otherwise specified. $C_\alpha$ RMSD, $C_\alpha$ RMSF, solvent accessible surface area (SASA), dihedral angles, residue-residue contacts, atom-atom contacts, and interatomic distances were calculated with *cpptraj* (20) Two residues were considered in contact with one another if at least one pair of heavy atoms were within 5 Å of one another. Atom-atom contact lifetimes were calculated using the previously described contact dataset and the *lifetimes* function of cpptraj with a fuzzy value of 2. A 5 Å cutoff was also used to identify atom-atom contacts. Set theory was used to assess the evolution of the BH – FH interface. For each snapshot in the simulation, the residue-residue pairs interacting across the BH – FH interface were grouped into sets. Then the Jaccard Index (JI), which ranges from [0,1] and is the ratio the intersection and union of two sets, was calculated to compare sets. High values of the JI indicate a stable set of residues at the BH – FH interface while low values indicate dynamic heterogeneity at the interface. Inter-helical angles for different contiguous α-helices in the tails were calculated using *cpptraj*. Residues forming helices were identified by selecting residues that adopted α-helical structure for at least 90% of the simulations. $C_\alpha$ atoms were chosen for alignment using a Low-Order-Value-Optimization strategy as described by Martínez and coworkers (21–23) and implemented in *mdlovofit* (*v20.0.0*) .(21) The myosin + ADP.Pi and myosin + dADP.Pi simulations were subsampled every 12 ns, non-protein atoms were removed, and then the two simulations were combined into a single ensemble. We calculated average structures for the aggregate ensemble using *cpptraj*. We used *mdlovofit* to identify the subset of $C_\alpha$ atoms with the smallest possible average RMSD to the averaged structures as a function of subset size (Φ). Protein images and movies were generated with *UCSF Chimera* (24) and *PyMOL* (25) .
